# Machine learning recognises senescence in glioblastoma and discovers senescence-inducing compounds

**DOI:** 10.1101/2024.04.03.587883

**Authors:** Lucy Martin, Anna Irving, Yossawat Suwanlikit, Gillian Morrison, Richard J.R. Elliott, Neil Carragher, Steven Pollard, Tamir Chandra

## Abstract

Senescence is a cell-intrinsic tumour suppressive response. A one-two-punch cancer treatment strategy aims to induce senescence in cancerous cells before removing them with a senolytic. It is important to accurately recognise senescent cells to investigate the feasibility of such a treatment strategy and identify compounds that induce senescence in cancer. We focus specifically on the terminal brain cancer glioblastoma, firstly identifying senescent glioblastoma cells with conventional stains, before training a machine learning model to distinguish senescent cells using only a DAPI nuclear stain. To demonstrate how our method can aid drug discovery, we apply our pipeline to existing glioblastoma high-throughput phenotypic drug screening imaging data to identify compounds that induce senescence in glioblastoma and verify these predictions experimentally.

**Author Summary:** Damaged cells can enter a senescent cell state, in which they do not divide, but continue to interact with the environment around them. A novel potential cancer treatment strategy is to make tumor cells senescent, before removing senescent cancer cells with a targeted drug. To investigate this treatment strategy in the brain cancer glioblastoma, it is important to be able to accurately recognise senescent glioblastoma cells. As identifying senescent cells is challenging, we create a machine learning pipeline which can detect senescent glioblastoma cells in imaging data. We show that by applying our method to existing data we can discover compounds that induce senescence in glioblastoma. We verify our predictions by testing the compounds experimentally.

## 1 Introduction

Senescent cells play a significant role in human ageing and disease. Characterised as a metabolically active state of proliferative arrest, senescence was first described in 1961 [1] and later identified as a cell-intrinsic tumour suppressor mechanism[2, 3]. More recently, pro-tumorigenic roles for senescent cells have been suggested, where they contribute towards an inflammatory tumour microenvironment (TME) [4, 5, 6].

Without a universal marker for senescent cells, a combination of common markers has been used for classification [7]. The absence of long-term BrdU incorporation is used to demonstrate proliferative arrest. Increased expression of p16 or p21 [8], a loss of laminB1 [9] and the presence of senescence-associated-*β*-galactosidase (SABG) [10, 11] have been used to identify senescent cells through imaging. Senescent cells and nuclei often display specific morphological phenotypes [12, 13, 14, 15].

Primary glioblastoma (GBM) is the most common and aggressive type of primary brain cancer in adults, with a median survival time of 15 months [16, 17]. The treatment for GBM is surgical resection followed by chemotherapy and radiotherapy [18]. However, even with treatment, cancer reoccurs. Both radiotherapy and chemotherapy have been found to induce senescence in GBM cells [19, 20], and although there is mounting evidence that senescence burden leads to poorer outcomes for GBM patients [21, 22], we currently do not understand the role of senescence in treatment. Furthermore, primary GBM tumours show a mutational spectrum consistent with senescence escape, with frequent mutations in the TERT promoter and CDKN2A, indicating that escape from senescence likely plays a role in the etiology of GBM [23].

Recently, a “one-two-punch” strategy for cancer treatment has gained popularity (Fig. 1a) [24, 25]. The treatment aims to induce senescence, specifically in tumour cells, before killing these cells with a senolytic. A one-two-punch strategy has the potential to not only be an effective treatment but also to reduce the likelihood of recurrence by preventing senescent cells from contributing towards a protumorigenic microenvironment [26]. Evidence suggests that a one-two-punch strategy may work in the brain, as senolytics have been shown to effectively remove senescent cells after radiation treatment [26].

**Figure 1:**
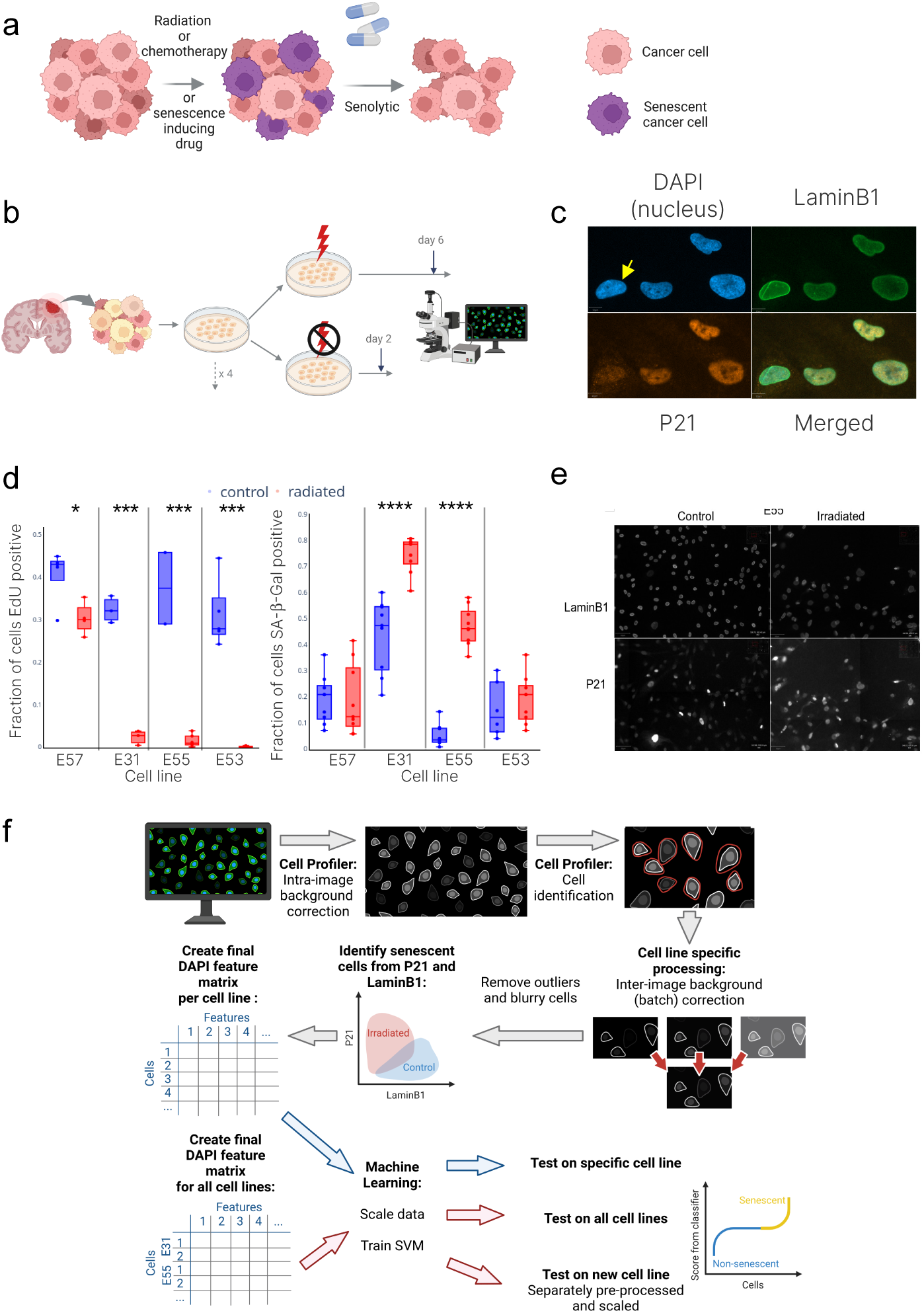
A pipeline to identify senescent glioblastoma cells. a) A “one-two-punch” strategy can drive cells into senescence before eliminating them. b) An outline of our experimental procedure: we induced senescence with radiation before IF staining the cells. c) An example of the p21, laminB1 and DAPI stain, the yellow arrow points to a non-senescent cell with lower p21 and higher laminB1 expression. d) The fraction of cells positive for SABG and EdU incorporation in the control and post-radiation. e) An example of the loss of laminB1 and gain of p21 post-radiation. f) An overview of our cell identification and machine learning pipeline.

Increasingly, a combination of high-throughput drug screening and machine learning is used to advance drug discovery [27, 28]. In vitro cell lines are treated with libraries of small-molecule compounds, and high-content imaging is used to automatically acquire images of cells after treatment [29]. Pipelines capable of analysing a large number of images search for compounds which lead to cell death or phenotypic change, by first performing in-depth image processing [30] and then applying machine learning algorithms [31]. These “hit” compounds are investigated further to determine if they can be used therapeutically.

Two recent papers have used DAPI and machine learning techniques to quantify senescence. The first used deep learning methods [12], and the second used feature extraction followed by random forest and tree-based classifiers [32]. Although these methods claim to generalise well across cell types and to be applicable in vivo, for the greatest accuracy, they must be trained on the cell type that they will be used on. Given the heterogeneity in the mutational spectrum and morphology of the GBM cell lines, we developed a GBM-specific senescence classifier using only features obtained from DAPI staining, enabling us to use existing imaging datasets to search for compounds which induce senescence in GBM.

Using cell labelling with multiple stains to identify senescence in GBM is a complicated, multi-step process that lacks clarity in results and reproducibility. A single method of senescence classification will ensure that senescent cells in vitro can be identified easily and in a cost-effective manner for high throughput screening, potentially aiding in the discovery of drugs that induce senescence in glioblastoma. In this paper, we identify senescent GBM cells in four patient-derived GBM cell lines using laminB1 and p21 stains to create a unique training set. We develop a novel GBM senescence classifier which can be applied to existing drug screening resources. As an example, we apply our pipeline to reanalyse existing image-based high-throughput drug screening data, identifying several compounds as senescence-inducing. Of these compounds, a significant fraction are glucocorticoids (GCs). While glucocorticoids are involved in GBM treatment, there are conflicting reports of whether they help or hinder tumour progression. Similarly, the mechanism of GC crosstalk with GBM remains poorly understood, such as whether their action is on the environment or the tumour cells. Our data indicates a direct interaction of GBM cells and GCs through the induction of senescence [33, 34].

## 2 Results

We used radiation (6Gy, x-ray) to induce senescence in four patient-derived glioblastoma cell lines (E55, E57, E31, and E53, see Table 1) before using an immunofluorescence (IF) stain for laminB1 (LMNB1), cyclin-dependent kinase inhibitor p21 (p21), and DAPI (Fig. 1b and c). We confirm senescence post-radiation with EdU incorporation and SABG staining in addition to laminB1 and p21 (Fig. 1d). We extracted over 300 quantitative features per cell relating to the p21, laminB1 and DAPI stain in both the irradiated and control cells using a CellProfiler image analysis pipeline (Fig. 1c and e), which is described in detail in the Methods (Section 4). Features quantify the size and shape of the nucleus in addition to the intensity of all three stains.

### Senescent glioblastoma cells can be characterised by the loss of laminB1 and the gain of p21

After pre-processing and normalising the data (Section 4, Methods), we sought to identify cells as senescent based on the increased p21 expression and loss of laminB1. LaminB1 is predominantly expressed in the nuclear envelope and is observed as a high-intensity ring around the nuclear perimeter (Fig. 2a), whereas p21 is expressed predominantly in the nucleus.

**Figure 2:**
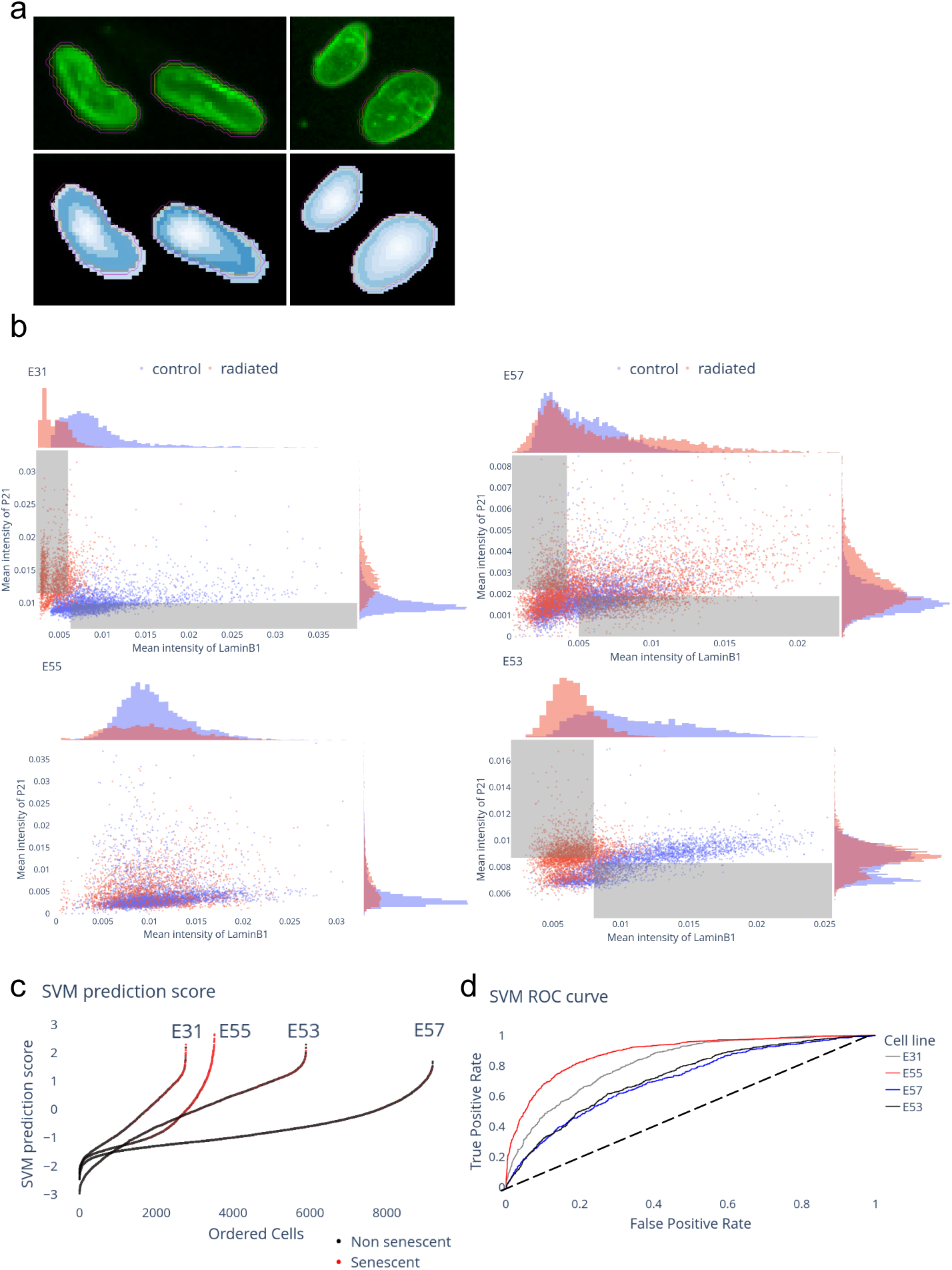
Identifying senescent glioblastoma cells from nuclear morphology. a) Upper panels show the laminB1 stain for control (left) and irradiated (right) E31 cells. Lower panels show the amount of stain in each segment of the nucleus. The orange line shows the nucleus identified by the DAPI stain, and the pink line is the expanded area used to identify the laminB1 stain. b) Identification of senescent and non-senescent cells based on the laminB1 and p21 stain. Control cells are blue and irradiated in red; grey boxes show the classification threshold. c) The predicted senescence score per cell for the test dataset, with cells coloured red if they were identified as senescent based on the levels of p21 and laminB1 and ordered by predicted senescence score. d) The ROC curve for the SVM trained using only the cells identified as senescent and non-senescent and applied to all cells.

Each cell line was processed independently as they expressed differing basal levels of p21 and laminB1. This difference in basal and post-radiation expression was unsurprising, as GBM is a highly heterogeneous cancer. The four cell lines were morphologically distinct, even by phase microscope imaging, where they could be easily distinguished under the microscope and had differing division rates.

In three of the four cell lines, we saw a loss of laminB1 and up-regulation of p21 in a subset of cells after radiation (Fig. 2b), with the most apparent distinction in cell line E31. We do not see clear changes in the quantitative values of intensity of laminB1 and p21 extracted from the CellProfiler pipeline for cell line E55 (Fig. 2a). However, EdU incorporation and SABG staining suggest that almost all E55 cells become senescent after radiation (Fig. 1d).

For cell lines E31, E53, and E57, we used a threshold in our metrics for p21 and laminB1 to select a subset of cells that showed low expression of laminB1 and high expression of p21 (Section 4.3.7); these were classified as senescent (Fig. 2b). In the same way, non-senescent cells were identified as cells with high laminB1 expression and low p21 expression (Fig. 2b). For E55, based on the SABG staining and EdU incorporation, we classified all radiated cells as senescent.

### Senescent glioblastoma cells can be identified with machine learning methods using only a DAPI stain

Using the subset of cells that we had identified to be senescent from the laminB1 and p21 stain, we trained several machine-learning models using only the features extracted from the DAPI staining of the cells (*∼* 100, features relating to the laminB1 and p21 stains were discarded). To account for uncertainty associated with the senescence classification, we chose methods that would also output the probability that a cell is senescent.

We used three supervised machine learning methods: a support vector machine (SVM), adaptive boosting (AdaBoost), and a boosted decision tree. Initially, we considered each cell line separately, training the classifier on a subset (50%, justified in Fig. S2b) of the data for each cell line and testing the classifier on the remaining cells. We found that all three models perform well across all four cell lines (Table S2, Table S3). To allow our model to be easily applied to feature data from other CellProfiler pipelines, we reduced the number of features used by the model to 30 features commonly extracted by most CellProfiler pipelines (Table S4); this did not adversely impact the performance of our models.

As we trained our classifiers with only a subset of senescent cells, those with the highest p21 and lowest laminB1 expression, we assume that we have underestimated the number of senescent cells in the training set. There will likely be a population of cells not initially labelled as senescent based on the intensity of laminB1 and p21 that are senescent and, therefore, have a senescent-like nuclear morphology. This was reflected in the large drop in precision (out of those cells predicted to be senescent, how many are senescent based on the laminB1 and p21 stain) when the models were tested on all cells.

All three classifiers return a score indicating the likelihood that a given cell is senescent, and the performance of each model when trained and tested on E31 is summarised in Table S3. For the remaining analysis, we used the SVM classification as it performed best in all metrics over all cell lines and returned a distribution of senescence scores with few outliers (Fig. 2c).

For all cell lines, cells classified as senescent by the laminB1 and p21 stain have a higher senescence prediction score in the test set (Fig. 2c). Of the three cell lines in which we could quantify a change in p21 and laminB1 expression, the SVM trained and tested on cell line E31 performs best (an AUC of 0.82, vs. 0.75, Fig. 2d). This is unsurprising given the large change in p21 and laminB1 post-irradiation in cell line E31, suggesting that E31 undergoes a more distinct senescence transition. Furthermore, we find that in a t-distributed stochastic neighbour embedding (t-SNE) reduction of the DAPI feature data for cell line E31, the location of cells with higher predicted senescence scores matches the location of cells classified as senescent by laminB1 and p21 (Fig. S1, a and b).

To test the cell line specificity of our model, we trained and tested an SVM on a mixture of cells from all four cell lines. We evaluated the overall performance of this more general model and found that the performance was worse (AUC of 0.69), as expected, due to patient heterogeneity. The performance of the SVM on each cell line is given in Table S2.

These results indicate that our classifier can accurately identify senescent GBM cells from their nuclear morphology.

### Comparison of nuclear features of senescent GBM cells to known features of senescence

Previous studies have identified nuclear changes in fibroblasts with senescence through feature extraction [13], and deep learning models [12]. In fibroblasts, cells become larger in area and show changes to the nuclear envelope, with one study also showing that senescent cells have a larger convexity (a ratio of context hull perimeter to perimeter, a measure of how jagged the nuclear membrane is). However, morphological changes are known to be cell line-dependent; considering GBM cells are mutated in several senescence pathways, we did not necessarily expect our cell lines to behave in the same way as karyotypically normal fibroblasts.

We used two algorithms to identify the importance of each feature in the SVM model trained on cell lines E31 and E57 (Fig. S3). First, using a permutation importance algorithm [35], we found that across the cell lines, the most important features are related to nuclear size and shape (e.g. “areashape compactness”, a measure distinguishing between nuclei that resemble filled circles, and irregular or irregularly stained nuclei) or the edge intensity of the DAPI stain (describing the nuclear envelope)(Fig. S3a). This suggests that we saw some of the morphological changes previously described in fibroblasts.

Using a game-theory-based approach (calculating SHAP values, SHapley Additive exPlanations [36]), we found that the three most important features were related to the intensity of the DAPI stain, not the nuclear size (Fig. S3b). However, we saw that cells with a larger nuclear extent and compactness and with a lower form factor and solidity are more likely to be senescent (Table S4), supporting the idea that senescent cells are more irregular or jagged in shape.

### Application to drug screening datasets to identify compounds inducing senescence

To find compounds that induce senescence in GBM cells, we applied our classification pipeline to the data generated in high-throughput drug screening experiments in which two of our four initial cell lines, E31 and E57, were used.

The cells were treated with compounds from two drug libraries, Targetmol (384 compounds, 4 concentrations) and LOPAC (1280 compounds, 2 concentrations), for 72 hours before the cells were fixed and stained with DAPI as part of a cell painting assay (Section 4).

We applied our machine learning pipeline to feature extraction data from the drug screening experiment, as raw images had already been processed with a CellProfiler pipeline. We calculated the mean senescence score for cells treated with each compound and the fraction of cells identified as senescent for each compound (Fig. 3a). Some compounds killed GBM cells, resulting in fewer live cells at the end of the treatment. For smaller total cell numbers, we expect to see a greater variance in the average senescence score per compound. Therefore, we used bootstrapping to calculate a cell number-dependent significance threshold (Fig. 3a green points, details in Section 4.4.3). Compounds that exceeded this threshold (Fig. 3a), both in average senescence score and the fraction of senescent cells for both cell lines, were classified as potential inducers of senescence. Compounds can be grouped into positive controls (genotoxic compounds known to induce senescence, such as etoposide), test compounds, negative controls (DMSO, the solvent used for all compounds), and cell-killing controls (paclitaxel (PAC), a microtubule-stabilizing agent that arrests cells in mitosis and can lead to cell death). We expected small concentrations of DMSO to neither reduce the number of cells (by causing cell death) nor induce senescence in cells, which was confirmed in our data (Fig. 3b, magenta points). The cell-killing control PAC killed glioblastoma cells and increased the senescence score (Fig. 3b, red points). Although evidence suggests that PAC kills glioblastoma cells, there is currently no evidence in the literature that PAC induces senescence or leads to morphological cell changes.

**Figure 3:**
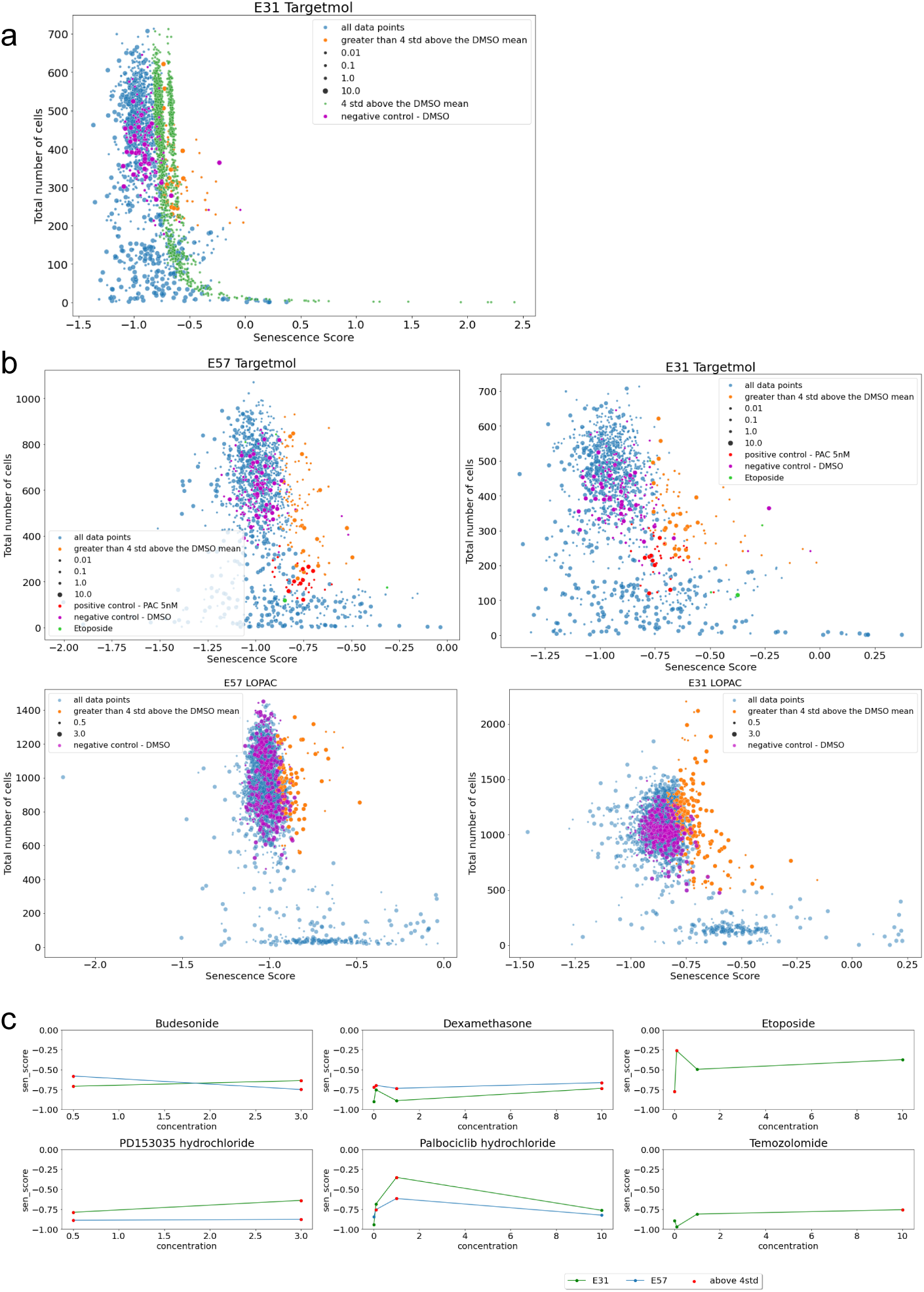
Using the machine learning model to identify senescent cells from drug screening. a) Average senescence score for compounds from the targetmol library, applied to cell line E31, showing the bootstrapped derived 4 standard deviations from the DMSO mean in green. b) Senescence scores with interesting compounds identified (orange), and DMSO controls, PAC, and Etoposide are highlighted. c) Senescence scores of compounds verified in the lab, with data points 4 standard deviations above the DMSO mean highlighted in red.

We chose to investigate compounds that caused a significant increase in the senescence score without killing large numbers of cells (Fig. 3a, orange points). Focusing on non-cytotoxic compounds, we identified senescence inducers that may be used as part of a one-two-punch treatment.

We identified approximately 20 candidates for senescence-inducing compounds in the cell lines E31 and E57; several were GCs (Table S1). However, it is also worth noting that some GCs seem to increase the number of glioblastoma cells in the drug-screening datasets. Of the four compounds in the LOPAC library that increased the cell number to above 1500 and induced significant levels of senescence in E31 (Fig. 3b, lower right plot, orange), three are GCs, suggesting that GCs may increase GBM cell proliferation (Fig. S5a).

To verify that the results are unaffected by small changes in the CellProfiler pipeline and, therefore, the feature extraction, we re-processed a selection of raw images taken as part of the drug screening experiments. We ran our CellProfiler pipeline on the images corresponding to two compounds of interest, and one DMSO control, extracting *∼* 100 DAPI features (not just the 30 features used in the simplified model), before determining the senescence score associated with each cell (Fig. S4c). We see significantly elevated levels of senescence in the compounds of interest compared to the DMSO control.

One GC of interest is dexamethasone, a compound often used in GBM treatment to reduce brain edema and inflammation and significantly improve patient quality of life. The effect of dexamethasone on GBM cells is an area of active research [33], with some studies suggesting that it may lead to increased cell proliferation, migration, and therapy resistance. Although no specific link has been made between dexamethasone and senescence in GBM cells (it has been found to induce senescence in lung epithelial cells [37]), it has been reported that dexamethasone induces p21 expression and inhibits apoptosis. If dexamethasone induces senescence in GBM cells, this may explain both the chemo- and radio-resistance and negative effects on survival rates if these senescent cells help to create a protumorigenic TME.

We found that both dexamethasone and dexamethasone acetate produced similar senescence scores in our two cell lines. Furthermore, other chemically similar compounds produced similar senescence scores, suggesting our pipeline worked as intended. To determine if our pipeline can identify senescence induction due to a range of chemically distinct compounds, we represented our compounds using the Simplified Molecular Input Line Entry System (SMILES). We performed dimensionality reduction of these chemical features with UMAP. We found that senescence-inducing compounds identified by our machine learning method were chemically diverse (Fig. S4b).

### Laboratory verification of senescence induction

To test the performance of our machine-learning model, we chose four of the compounds that were predicted to be senescence-inducing to test in the lab (dexamethasone, PD153035, palbociclib hydroxide, and budesonide), alongside two positive controls (temozolomide and etoposide). Etoposide, a topoisomerase II inhibitor, is commonly used to induce senescence in many cell lines [32], and significant evidence now shows that the current standard of care chemotherapy drug to treat glioblastoma, temozolomide, induces senescence in GBM cells [20]. The concentrations of these compounds predicted to give the maximum senescence induction were used in the experiment (Table 2, Fig. S5b). To replicate the conditions of the drug screening experiments, compounds were applied for 72 before cells were fixed and stained.

For simplicity, we used only cell line E31 and stained for p21 to indicate senescence. In all six of these compounds, we see an increase in p21 intensity compared to the control cells, which were treated only with DMSO (Fig. 4a). This increase was significant for all compounds. However, the effect size differed between compounds, with the largest change in etoposide-treated cells (a 2.52 fold-change in p21 expression, compared to a 1.52 fold-change in dexamethasone). Using a threshold (arbitrary) in p21 expression to determine senescence, we found that all compounds also showed a significant increase in the number of senescent cells observed (Fig. 4b). The largest increase in the mean intensity of p21 per cell and the fraction of senescent cells was in etoposide, as predicted by our model (Fig. 3c).

**Figure 4:**
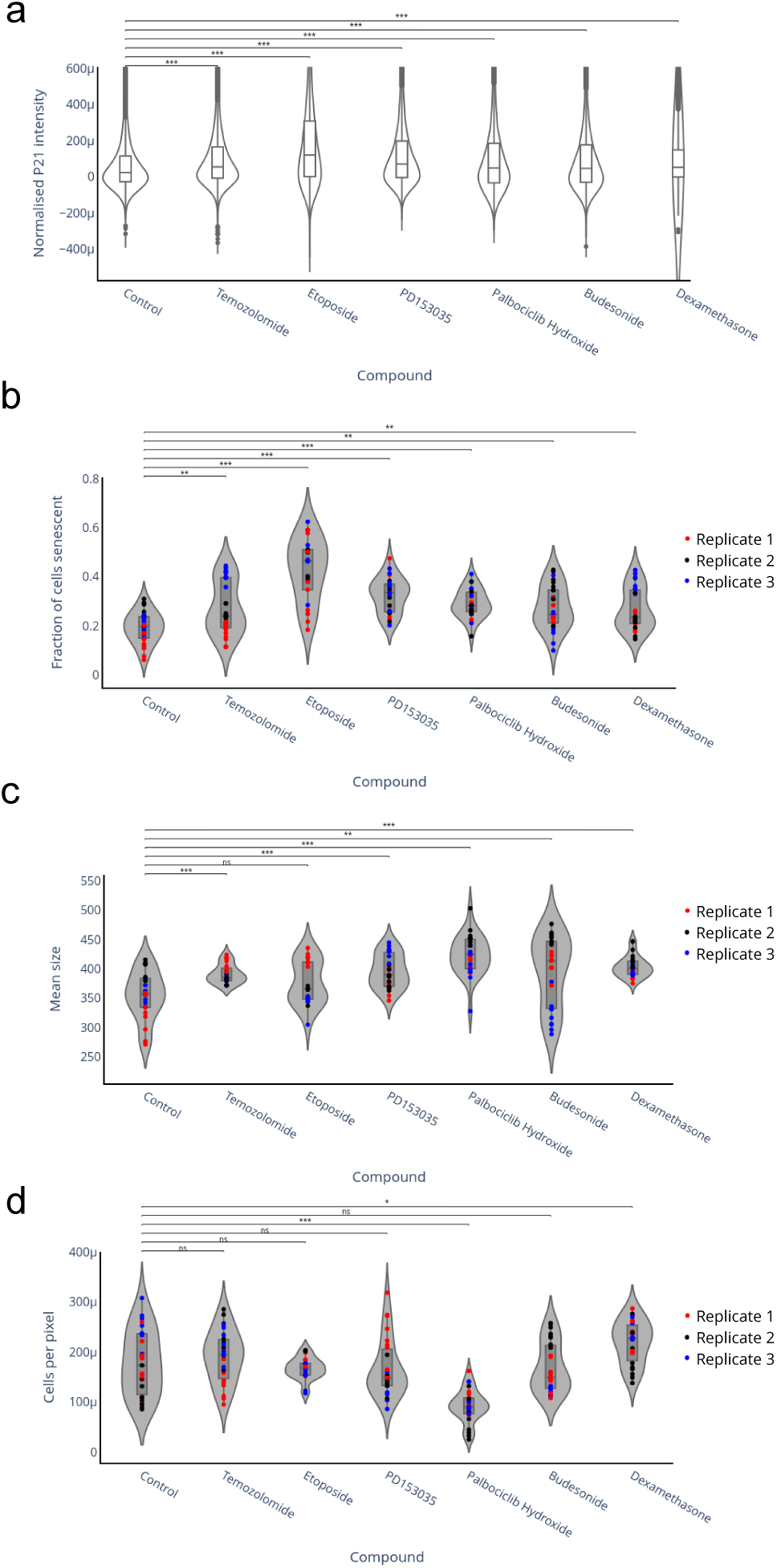
Laboratory testing of potentially senescence-inducing compounds. a) The normalised p21 intensity in each cell for each applied compound. b) The fraction of senescence in each replicate (denoted by the colour of points) for each applied compound. c) The mean size of cells in each replicate (denoted by the colour of points) for each applied compound. d) The number of cells per pixel (per unit area) in each replicate (denoted by the colour of points)

All compounds led to a small increase in cell size (Fig. 4c). However, the changes in cell size observed did not correlate with the changes in p21 expression, supporting the conclusion that simple measures of morphological change are insufficient to predict senescence. Furthermore, only one of the compounds tested, dexamethasone, appeared to cause increased proliferation of the GBM cells (Fig. 4d), and this increase was small (fold-change of 1.23).

## 3 Discussion

Glioblastoma is a cancer of unmet need. Although understanding of this cancer has improved in the last decade, this has not translated into new therapeutic options. Senescence is heavily implicated in GBM progression, with several recent studies showing that a higher senescence burden before treatment can lead to poorer patient outcomes and that chemotherapy and radiotherapy lead to therapy-induced senescence in GBM. Furthermore, nearly all GBMs are mutated in pathways associated with senescence, indicating that although GBM cells can become senescent, the senescent phenotype is likely to differ from the senescence observed in healthy cells.

With an increased understanding of senescence in GBM, it may be possible to leverage therapy-induced senescence as part of a one-two-punch strategy, first inducing senescence specifically in GBM cells before clearing these cells with a senolytic. To do this, we need an effective way of identifying senescent GBM cells and drugs that induce senescence in GBM.

We have created a dataset containing images of four GBM patient-derived cell lines with and without radiation treatment. We identify senescent cells through immunocytochemistry p21 and laminB1 staining and develop a machine-learning pipeline to identify senescent GBM cells based only on a DAPI nuclear stain. Applying our pipeline to high-throughput drug screening data, we identified 20 compounds that we predict induce senescence in GBM cells. Our pipeline can be applied to any GBM in vitro imaging data with a DAPI stain, allowing existing high throughput drug screening data to be used to its full potential to explore the senescent phenotype.

For example, our machine-learning model identifies dexamethasone (and several other GCs) as a compound that may cause senescence in GBM cells. However, the high-throughput drug screening data also suggests that GCs may lead to increased proliferation in some GBM cell lines.

We tested four of our hit compounds in vitro. We found that all compounds increased p21 expression in cell line E31. While the increase was significant for all compounds, the effect size varied, with the positive control etoposide leading to the largest change in p21 expression. Furthermore, only one of the compounds tested (dexamethasone) led to a small increase in cell proliferation, suggesting that senescence induction is not simply a result of increased proliferation and overcrowding and that GCs do not cause a significant increase in proliferation in this cell line.

This study has several limitations. First, we only induce senescence through a single mechanism, radiation, and the senescence phenotype is known to vary between induction mechanisms. Second, we tested our classifier using a dataset in which GBM cells were treated with compounds for 72 hours before cells were fixed and stained. This may not be sufficient time for the senescence phenotype to fully develop.

Furthermore, additional work will be needed to test hit compounds before they can be used in a one-two-punch treatment strategy. For example, showing that compounds induce senescence selectively in GBM cells, they do not affect healthy brain cells, and that the senescent cells can be removed with a senolytic. Investigation of the mechanism of action of hit compounds may help identify which cells are vulnerable to senescence induction.

In summary, our findings demonstrate the potential of machine learning classifiers to be applied to determine distinct cellular states and responses to therapy to help in new drug discovery efforts for GBM.

## 4 Methods

### 4.1 Key Resources Table

**Table 1:** Key Resources Table.

| REAGENT OR RESOURCE | SOURCE | IDENTIFIER |
| --- | --- | --- |
| Antibodies |  |  |
| Anti-LaminB1 antibody | abcam | Cat#ab16048 |
| Goat Anti-Rabbit IgG (H+L), highly cross-adsorbed, Hilyte Fluor™ 488-labeled | ANASPEC | Cat#AS-61056-05-H488 |
| Anti-p21WAF1/Cip1 antibody, Mouse monoclonal | Sigma-Aldrich | Cat#P1484 |
| Goat Anti-Mouse IgG (H+L), highly cross-adsorbed, Hilyte Fluor™ 555-labeled | ANASPEC | Cat#AS-61057-05-H555 |
| Chemicals, peptides, and recombinant proteins |  |  |
| Dulbecco's modified eagle medium (DMEM) with Ham's F-12 | Sigma-Aldrich | Cat#D8437 |
| D-(+)-Glucose solution | Sigma-Aldrich | Cat#G8644 |
| MEM Non-Essential Amino Acids Solution (100X) | Gibco | Cat#11140-035 |
| Penicillin-Streptomycin (10,000 U/mL) | Gibco | Cat#15140-122 |
| Bovine Albumin Fraction V (7.5% solution) | Gibco | Cat#15260-037 |
| 2-Mercaptoethanol (50 mM) | Gibco | Cat#31350-010 |
| B-27™ Supplement (50X), serum free | Gibco | Cat#17504-044 |
| N-2 Supplement (100X) | Gibco | Cat#17502-048 |
| Recombinant Murine EGF | PeproTech | Cat#315-09 |
| Recombinant Human FGF-basic (154 a.a.) | PeproTech | Cat#100-18B |
| 3-D Culture Matrix Laminin I | Cultrex | Cat#3446-005-01 |
| Accutase® Cell Detachment Solution | BioLegend | Cat#424201 |
| Dimethyl sulfoxide (DMSO) | Sigma-Aldrich | Cat#D2650 |
| Etoposide 10 mM (in 1mL DMSO) | Apexbio | Cat#A1971-APE |
| Palbociclib Hydrochloride | Cambridge LKT Labs | Cat#827022-32-2 |
| Dexamethasone (Dex) | Cell guidance systems | Cat#50-02-2 |
| Temozolomide | Cayman chemicals | Cat#85622-93-1 |
| PD153035 (Hydrochloride) | MedChem Express | Cat#880813-42-3 |
| Critical commercial assays |  |  |
| Click-iT™ EdU Alexa Fluor™ 647 HCS Assay | Invitrogen | Cat#C10356 |
| Deposited data |  |  |
| Experimental models: Cell lines |  |  |
| Human Glioma Stem Cells: E31, E53, E55, E57 | Steven Pollard Lab, Centre for Regenerative Medicine, Edinburgh, UK | N/A |
| Software and algorithms |  |  |
| CellProfiler v4.2.4 | Open Source | www.cellprofiler.org |
| Python v3.9 |  |  |
| Other |  |  |
| RX-650 X-Ray Unit | Faxitron | Cat#43855D |

**Table 2:** Compound concentrations.

| COMPOUND | CONCENTRATION USED |
| --- | --- |
| Etoposide | 0.1 μM |
| Palbociclib hydroxide | 0.1 μM |
| Dexamethasone | 0.1 μM |
| Temozolomide | 10 μM |
| PD153035 hydrochloride | 0.5 μM |
| Budesonide | 0.5 microM |

### 4.2 Experimental Model and Subject Details

The four glioblastoma cell lines were from The Pollard Lab at the Centre for Regenerative Medicine, University of Edinburgh. Patient-derived GSC lines were obtained from the Glioma Cellular Genetics Resource (https://gcgr.github.io), funded by a Cancer Research UK Accelerator Award (A21922).

#### 4.2.1 Cell Culture

Cells were cultured in a complete media of DMEM/HAMS-F12 (Sigma-Aldrich) supplemented with Glucose solution (Sigma-Aldrich), MEM NEAA 100x (Gibco), Penicillin-Streptomycin (Gibco), Bovine Serum Albumin Solution 7.5% (Gibco), 2-Mercaptoethanol (Gibco), B-27 supplement 50x (Gibco), N-2 supplement 100x (Gibco), human FGF (to a final concentration of 10 ng/ml)(Peprotech), murine EGF (to a final concentration of 10 ng/ml)(Peprotech), and laminin (to a final concentration of 2 ng/ml)(Culturex). For splitting, passaging, freezing, and thawing, a wash media of DMEM/HAMS-F12 (Sigma) supplemented with Glucose (Sigma) and Penicillin-Streptomycin (Gibco) was used.

The cells were grown in a complete media on CoStar Tissue Culture 25 cm^2^ (T25) plates and kept at 37 °C in 5% CO_2_-humidified incubators. Media was changed every 5-10 days if necessary, and cells were split or passaged every 5-10 days, depending on the growth rate of the cell line. When passaging or splitting, cells were removed from their plates with Accutase solution (BioLegend). Cells were split 1:4 or 1:6 depending on the cell line.

For temporary storage throughout the experiment, cells were suspended in a solution of 10% DMSO (Sigma-Aldrich) in wash media and kept at −80 °C in 1 ml aliquots. Recovery times from freezing varied by cell line.

For longer-term storage, cells were kept in liquid nitrogen.

### 4.3 Method Details

#### 4.3.1 Irradiation

Cells were irradiated with 6 Gray (Gy) using an RX-650 Faxitron X-ray unit. All cells were transported to the radiation unit, and non-radiated controls were kept out of the incubator for the same period of time as radiated cells. One day prior to irradiation, cells from a T25 plate at around 80-90% confluency were passaged and used to seed glass coverslips in 12-well plates at a dilution of 1:6. The media was changed 4 days after radiation.

#### 4.3.2 Immunofluorescence Staining

Firstly, cells were fixed and permeabilised using 3.7% formaldehyde followed by 0.5% Triton X-100. Washed cells were then incubated with the primary antibody against laminB1 for 45 minutes in the dark at room temperature, followed by the secondary antibody under the same incubation conditions. This was repeated for the primary and secondary antibodies against p21. Between incubations, cells were washed three times with phosphate-buffered saline (PBS) with 0.1 % tween-20 (PBST). Cells were mounted using a mounting medium with DAPI (Vectashield).

#### 4.3.3 Compound testing

Six compounds were tested to determine if they induced senescence as predicted by our classifier; four of these were strong hits from the classifier (dexamethasone, PD153035, palbociclib hydroxide, budesonide), and two were positive controls (temozolomide and etoposide). DMSO was used as a negative control.

One day prior to the application of compounds, cells from a T25 plate at around 80-90% confluency were passaged and used to seed glass coverslips in 12-well plates at a dilution of 1:6. The compounds were applied to the cells for 72 hours (concentrations used are described in table 2, concentrations were reached through serial dilution in DMSO), after which the cells were fixed and stained. Staining for p21 was performed as described in section 4.3.2.

#### 4.3.4 Fluorescence Microscopy

Imaging was performed using an axioscan fluorescence microscope.

#### 4.3.5 High Content Feature Extraction

A CellProfiler [38] pipeline was used to identify cells and quantify DAPI, laminB1 and p21 staining from fluorescent images. DAPI staining was used to create an object set of nuclei, which could be used to extract measurements across the DAPI, laminB1 and p21 image sets.

Firstly, an illumination correction was carried out to remove uneven illumination patterns in images using a median filter followed by a division function. Corrected images were termed CorrDAPI, CorrLaminB1, and CorrP21.

Primary object identification was carried out using a manual threshold that varied by cell line from 0.001 to 0.003. To improve the consistency of the object identification module settings across images with varying background intensities, the mean intensity of CorrDAPI was subtracted from the image, and the output image was used for object identification. The objects identified were labelled as NucleiObject.

NucleiObject was used to extract the nuclei’s size and shape features and per-object measurements of intensity, intensity distribution, texture, and granularity from CorrDAPI and CorrP21.

Intensity measurements for CorrLaminB1 using NucleiObject failed to capture the characteristic ring of laminB1 around the edge of the nucleus. To correct this, a second object set (DilatedNuclei 1) was created by dilating NucleiObject with a size of 1. The DilatedNuclei 1 object set was used to extract the same measurements from CorrLaminB1.

Masks of DilatedNuclei 1 on CorrDAPI, CorrLaminB1 and CorrP21 were used to measure the background intensity per image.

The pipeline’s resulting output was a series of background intensity measurements per image and over 300 per-object measurements for DAPI, laminB1, and p21 staining, exported in a CSV file.

#### 4.3.6 Data Processing

Data processing was performed in Python. Measurements from the CellProfiler pipeline were imported, and metadata and cell positional data were removed.

Features related to the intensity of DAPI, p21, or laminB1 stains were rescaled using the background intensity levels of the image as a whole. For each cell, the mean background intensity was subtracted from intensity features.

Objects (cells) that were outliers (above the 95% quantile, or below the 5% quantile) in more than 23% of features were removed. 23% was chosen using the “elbow” in a histogram of the number of outlying features per cell.

To reduce the number of features per cell from over 300, only features that contained a large amount of variance were kept.

#### 4.3.7 Identification of Senescent Cells from laminB1 and p21

Cells above a manually derived threshold of mean p21 intensity and below a manually derived threshold for mean laminB1 intensity are classified as senescent. Conversely, those below a threshold in p21 and above a threshold in laminB1 are classified as non-senescent (Fig. 2c).

#### 4.3.8 Compound testing analysis

After the experimental procedure described in Section 4.3.3, slides were images with the axioscan microscope. Nine images were taken per slide. Images were processed, and features were extracted using the CellProfiler pipeline described in Section 4.3.5. Results were normalised to account for variation in p21 intensity between the control (DMSO) slides so that the mode p21 intensity in each slide was 0. This normalisation relies on the assumption that there are significantly more non-senescent cells than senescent cells in each slide. This assumption is supported by the data shown in Figure S4a.

#### 4.3.9 Classifying Cells from DAPI Stain

To identify cells as senescent based only on the DAPI stain, we created a feature matrix for each cell containing only features extracted from the DAPI stain. Each cell is labelled either senescent, not senescent, or unclassified, based on the thresholds described in Section 4.3.7 and shown in figure 2c.

We investigated three classification models (SVM, AdaBoost, and a boosted decision tree) from scikit-learn [39], training each model on only the senescent and non-senescent populations before testing it both on the remaining senescent and non-senescent cells and all the remaining cells, including those which were not classified as very senescent or non-senescent based on the laminB1 and p21 stains (Table S3). We found that the SVM model performed best over a range of metrics and outputted a continuous range of senescence prediction scores with few outliers. Therefore, we used the SVM model in further analyses (Table S2). As the machine learning pipeline will be applied to other datasets (from images taken with different microscopes and potentially different magnifications), we chose to normalise all data with respect to the control cells (un-radiated), as we expect a small number of senescent cells in vitro [21](confirmed through SABG staining and EdU incorporation). The scikit-learn standard scaler [40] was trained on the control cells only (removing the mean and scaling to unit variance), for both the training and test data, before applying the scaling to both the control and treated (whether with radiation or a compound) cells. This ensured that the model could be trained on one dataset and applied to another and that the fraction of cells determined to be senescent by the model was accurate, not relative.

#### 4.3.10 Applying Machine Learning to Drug Discovery Data

The machine learning classification pipeline described above was applied to data from high-throughput drug screening experiments performed by Richard J.R. Elliott from Professor Neil Carragher’s Drug Discovery programme at the Institute of Genetics and Cancer, University of Edinburgh. As part of the drug screening, the cell lines E31 and E57 were fixed and stained with a cell painting assay, following treatment for 72 hours with drugs from two different drug libraries, targetmol (330 compounds, four concentrations: 10, 1.0, 0.1, and 0.01 micromolar) and LOPAC (1280 compounds, two concentrations). The cell painting assay included a DAPI nuclear stain. All images were acquired with an ImageXpress-Confocal high-content screening platform integrated with PAA plate handling robotics.

The resulting images were processed with a CellProfiler pipeline created by the Carragher lab. From these high-throughput screening experiments, we received a matrix containing CellProfiler features describing each cell’s DAPI nuclear stain.

DMSO was used as a negative control for both the Targetmol and LOPAC libraries, with two DMSO wells per row on each 384-well plate for the Targetmol library and one DMSO well per row on each 384-well plate for the LOPAC library. In addition, the Targetmol library used Paclitaxel (PAC) as a positive control, as it is known to kill glioblastoma cells.

To apply our pipeline to data produced from a different CellProfiler pipeline, we limited the features in our SVM model to those that also appear in the drug screening pipeline (*∼*30 features). This did not impact the performance of our model.

Our classification pipeline outputted senescence scores per cell, the fraction of senescent cells per well, the number of cells per well, and standard deviations for both the senescence score and the fraction of senescent cells per well.

#### 4.3.11 Identifying Interesting Compounds

From this output, compounds of interest were selected as compounds that induced a significant senescence response in cells. Significance was defined as greater than 4 standard deviations above the mean of DMSO controls. The compounds selected induced a senescence response in both cell lines (E31 and E57) through a significant increase in the mean senescence score and the fraction of senescent cells per well. Only wells with over 200 cells (targetmol) or 500 cells (LOPAC) remaining after treatment were selected to avoid choosing compounds that killed large numbers of cells, as this may induce senescence in the remaining cells.

DMSO control wells were included per plate in the experiments, so compoundtreated wells were compared to the DMSO controls of the same plate. Bootstrapping was carried out to eliminate bias from sample size in wells containing fewer cells (as described in Section 4.4.3).

#### 4.3.12 SABG Staining

Cells were stained with an X-gal solution which was left on for 19 hours/22 hours at 37 °C. The solution contained 90% PBS, 5% 20X KC, and 5% X-gal (ThermoScientific). 800 µl of X-gal solution was added per well of a 12-well plate. Prior to staining, cells were fixed with a 0.5% glutaraldehyde solution made using 25% glutaraldehyde stock (Sigma) diluted in PBS and left on cells for 12 minutes. After removing the X-gal solution, cells were kept in the dark at 4 °C.

#### 4.3.13 Bright-field Microscopy

Wells were imaged using a bright-field microscope. Three images were taken randomly per well of a 12-well plate, and all were taken in the same session.

### 4.4 Quantification and Statistical Analysis

#### 4.4.1 Cell Number

Cell numbers were quantified using CellProfiler’s primary object identification module for fluorescence and bright-field microscopy images (for full CellProfiler pipelines, see Data and Software Availability).

A manual threshold was selected for fluorescence microscopy images to identify cells from images in the DAPI channel. Different thresholds were set for different cell lines to account for differences in DAPI staining intensity. Prior to object identification, images were corrected for variations in background illumination, and the mean image background intensity was subtracted from the overall image to make identification more reliable across images.

For bright-field images, the manual threshold and size parameters were adjusted between cell lines to account for morphological differences. The original image was processed prior to the identify primary objects module to enhance cell shapes and increase the contrast between the cells and their background.

#### 4.4.2 Quantification of Senescence using SABG

CellProfiler was used to quantify blue X-gal staining from bright-field cell images. After background correction, we used the module unmix colours to extract blue shades from the original image. Unmix colours outputted a grayscale image where the highest intensity areas of the image reflected the areas of the input image with the most blue. This was quantified using primary object identification, with a manual threshold consistent across images and cell lines, identifying areas of stain within images that could be related to previously identified cells.

#### 4.4.3 Bootstrapping

To identify interesting compounds from the drug screening experiments described above, we set a significance threshold for senescence score and fraction senescence at four standard deviations above the control mean. Wells with fewer cells showed greater variance in mean senescence scores and the fraction of cells that were senescent. Because of this, we could not use a single standard deviation value to accurately reflect the significance of mean values from wells with smaller cell populations.

To account for this sample size effect on standard deviation, we applied bootstrapping with replacement to assign an expected standard deviation value per well, given the number of cells. In the original experiments, DMSO controls were included in each plate. For each well, we added 4 bootstrapped standard deviations to the mean of DMSO wells in the appropriate plate. This method was used for both senescence scores and predicted fraction of senescence and used to determine which wells fell above this significance threshold.

#### 4.4.4 Important feature identification

We used two algorithms, permutation feature importance and Shapley values, to identify important features in the SVM model.

We use the sklearn permutation feature importance algorithm [35], applied to the training data (50% of all data per cell line. Fig. S3a). Feature scores are randomly shuffled, and the model is reevaluated to determine which features impact the goodness of fit most. A caveat of this algorithm is that misleading values may be returned for highly correlated features.

We used the SHAP python package to calculate Shapley values for our model (Fig. S3b) [36]. This method is based on game theory, where features become players that can join or not join the game (model). If a feature has positive SHAP values for higher values of the feature, then higher values of that feature mean that a cell is more likely to be senescent.

### 4.5 Statistical significance

In Figure 4, statistical significance was calculated using a Mann–Whitney U test from the Python scipy.stats package.

## 4.6 Data and Software Availability

The CellProfiler pipeline and Python code used in this manuscript are available at https://github.com/lkmartin90/Image_ML_for_senescence.

## Contributions

L.M. and T.C conceived and supervised the study. L.M., A.I, and T.C wrote the manuscript. G.M. derived the GBM cell lines. S.P. curated the patient-derived cell lines and gave access to the cell lines. L.M. cultured and imaged the GBM cell lines, and developed the machine learning method. A.I developed the CellProfiler pipeline and image analysis. Y.S. and L.M performed the hit compound testing experiments. R.E and N.C. performed the high-throughput drug screening experiments and the CellProfiler processing of that data.

## Acknowledgments

L.M is a cross-disciplinary research fellow supported by funding from the CRUK Brain Tumour Centre of Excellence Award (C157/A27589). N.C and R.E. acknowledge funding support from Cancer Research UK and the Brain Tumour Charity (grant REF: C42454/A28596). Patient-derived GSC lines were obtained from the Glioma Cellular Genetics Resource (https://gcgr.github.io), funded by a Cancer Research UK Accelerator Award (A21922). We thank all members of the Chandra lab and the Advanced Imaging Resource (AIR), at the Institute of Genetics and Cancer, Edinburgh, for their input.

## Supplementary

**Table S1:** Compounds identified as senescence inducing.

| Compound Name | Drug Class |
| --- | --- |
| CB 1954 | Alkylating agent |
| Chloroambucil | Alkylating agent |
| Doxorubicin hydrochloride | Anthracycline |
| Selinexor | Anti-cancer agent |
| Teniposide | Anti-cancer agent |
| Beclomethasone | Corticosteroid |
| Budesonide | Corticosteroid |
| Dexamethasone acetate | Corticosteroid |
| Hydrocortisone butyrate | Corticosteroid |
| Nestorone | Corticosteroid |
| Triamcinolone | Corticosteroid |
| PD153035 hydrochloride | Kinase inhibitor |
| Palbociclib hydrochloride | Kinase inhibitor |
| CP466722 | Kinase inhibitor |
| Palbociclib isethionate | Kinase inhibitor |
| Cyclocytidine hydrochloride | Nucleoside analog |
| Cytarabine | Nucleoside analog |
| Talazoparib | PARP inhibitor |
| ClProtoporphyrin IX (PpIX) disodium | Porphyrin-based salt |
| 1-(4-Chlorobenzyl)-5-methoxy-2-methylindole-3-acetic acid | Small molecule |
| Wiskostatin | Small molecule |

**Table S2:** Metrics describing the performance of the SVM on all cell lines. “Subset”, is used when models were tested on only cells identified as senescent or non-senescent from the p21 and laminB1 stain.

| Cell line trained on | Cell lines tested on | Tested on (all or subset) | Accuracy | Precision | Recall |
| --- | --- | --- | --- | --- | --- |
| E31 | E31 | subset | 0.79 | 0.79 | 0.7 |
| E31 | E31 | all | 0.76 | 0.35 | 0.71 |
| E57 | E57 | subset | 0.82 | 0.79 | 0.55 |
| E57 | E57 | all | 0.85 | 0.15 | 0.44 |
| E55 | E55 | subset | 0.83 | 0.79 | 0.61 |
| E55 | E55 | all | 0.87 | 0.84 | 0.67 |
| E53 | E53 | subset | 0.82 | 0.86 | 0.86 |
| E53 | E53 | all | 0.61 | 0.22 | 0.79 |
| All | All | all | 0.77 | 0.75 | 0.58 |
| All | All | subset | 0.78 | 0.27 | 0.41 |
| All | E31 | subset | 0.77 | 0.34 | 0.61 |
| All | E55 | subset | 0.77 | 0.76 | 0.31 |
| All | E53 | subset | 0.61 | 0.19 | 0.63 |
| All | E57 | subset | 0.87 | 0.13 | 0.31 |

**Table S3:** Metrics describing the performance of the three tested models on cell line E31. “Subset”, is used when models were tested on only cells identified as senescent or non-senescent from the p21 and laminB1 stain.

| Model | Model details | Tested on<br>(all or subset) | Accuracy | Precision | Recall |
| --- | --- | --- | --- | --- | --- |
| SVM | Kernel = ‘rbf’ | subset | 0.78 | 0.78 | 0.71 |
| SVM | Kernel = ‘rbf’ | all | 0.75 | 0.46 | 0.7 |
| Ada Boost | n_estimators=100 | subset | 0.76 | 0.69 | 0.85 |
| Ada Boost | n_estimators=100 | all | 0.74 | 0.45 | 0.73 |
| Gradient boost | n_estimators=200,<br>learning_rate=1.0,<br>max_depth = 5 | subset | 0.75 | 0.7 | 0.75 |
| Gradient boost | n_estimators=200,<br>learning_rate=1.0,<br>max_depth = 5 | all | 0.77 | 0.5 | 0.72 |

**Figure S1:**
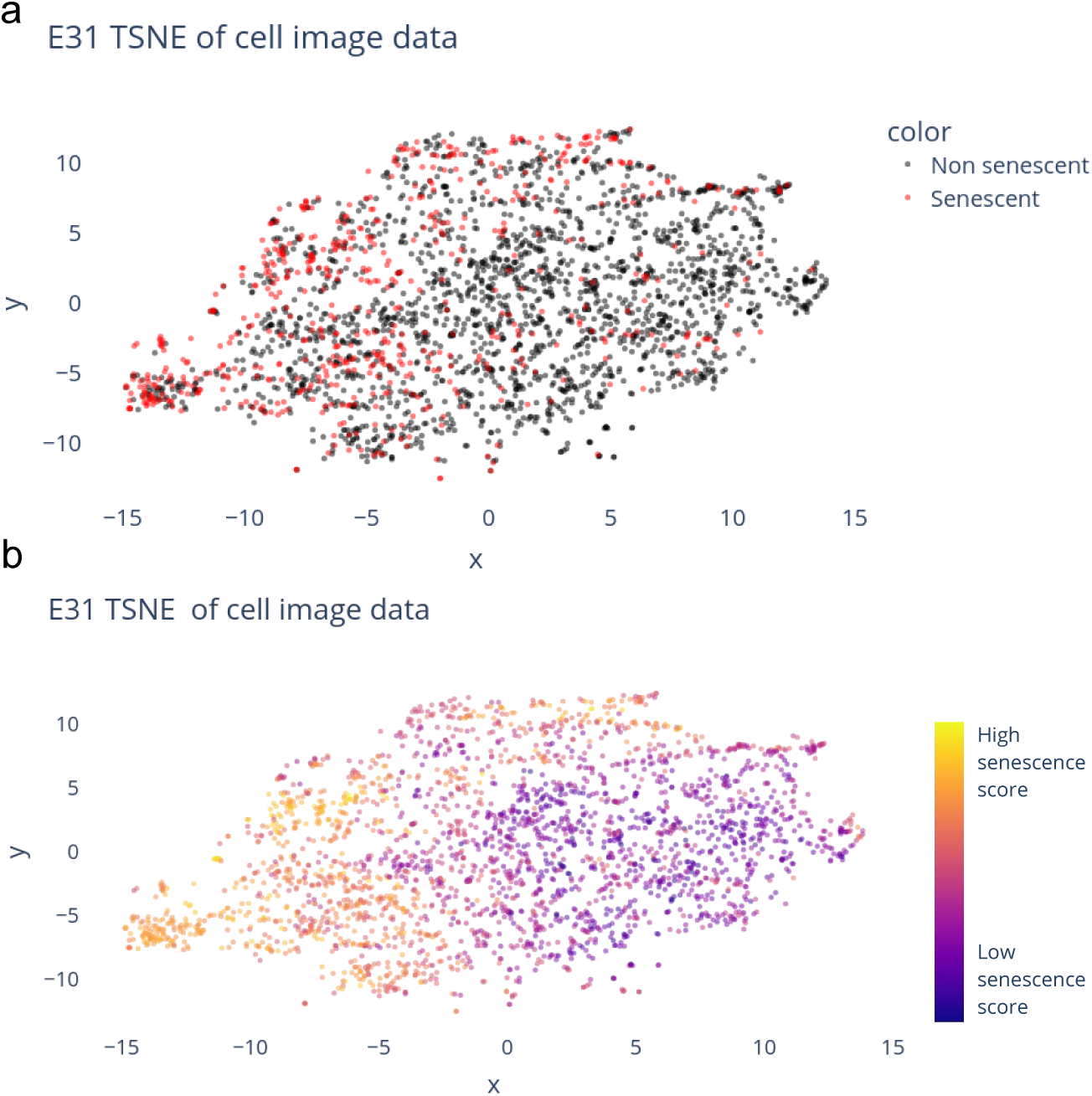
a) 2D TSNE reduction of the DAPI features for each E31 cell (both irradiated and control), coloured by whether the cells were identified as senescent based on the laminB1 and P21 stain (Fig. 2c). Yellow points are cells that were identified as very senescent-like, and blue points are cells that weren’t. b) The same TSNE reduction as in (a), coloured by the senescence score from the SVM model.

**Figure S2:**
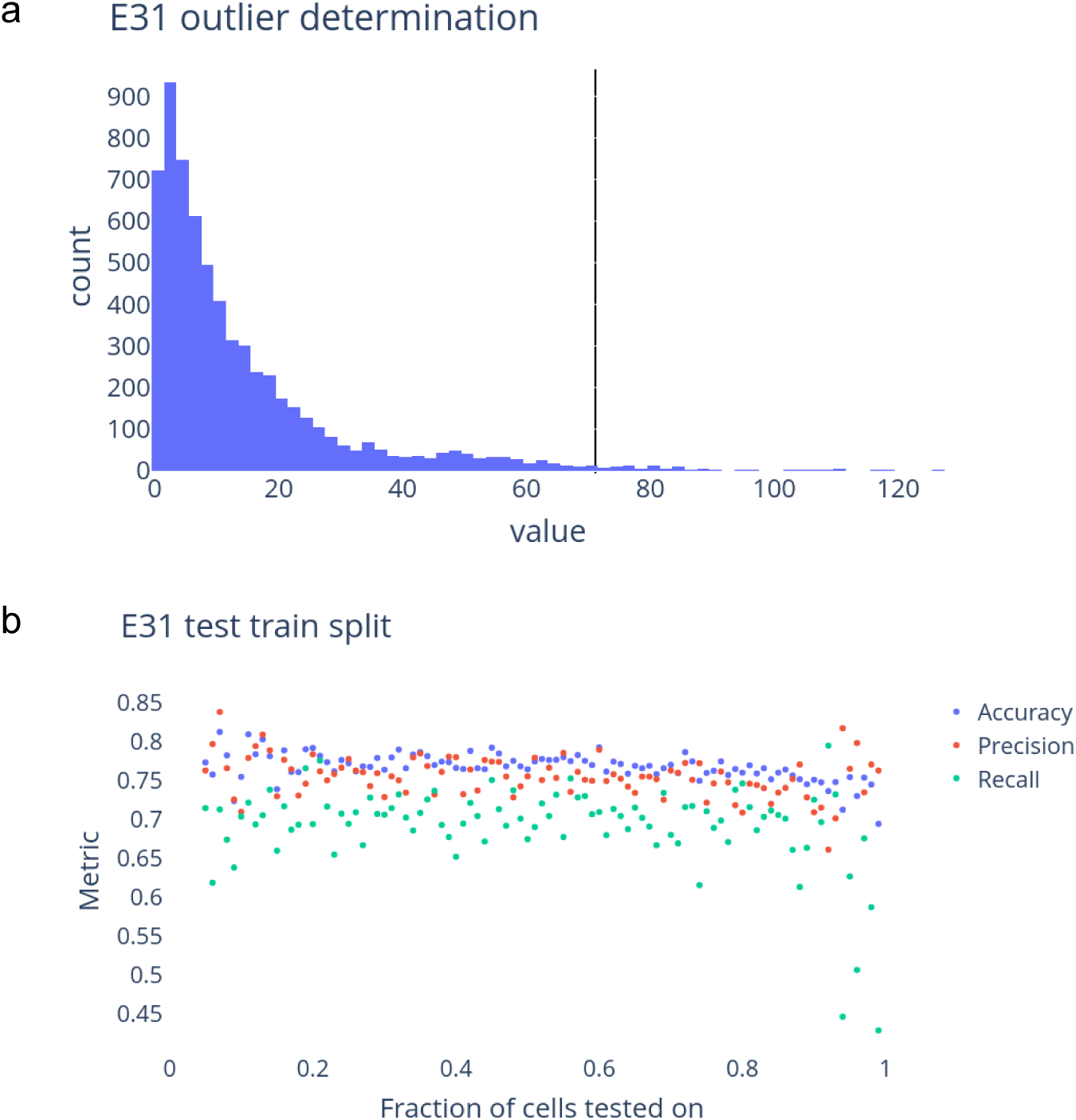
a) Histogram showing the outlier metric for removing outlier cells, for cell line E31. The vertical black line shows the threshold above which cells were discarded. b) Performance of the SVM on cell line E31 as a function of train/test set size.

**Figure S3:**
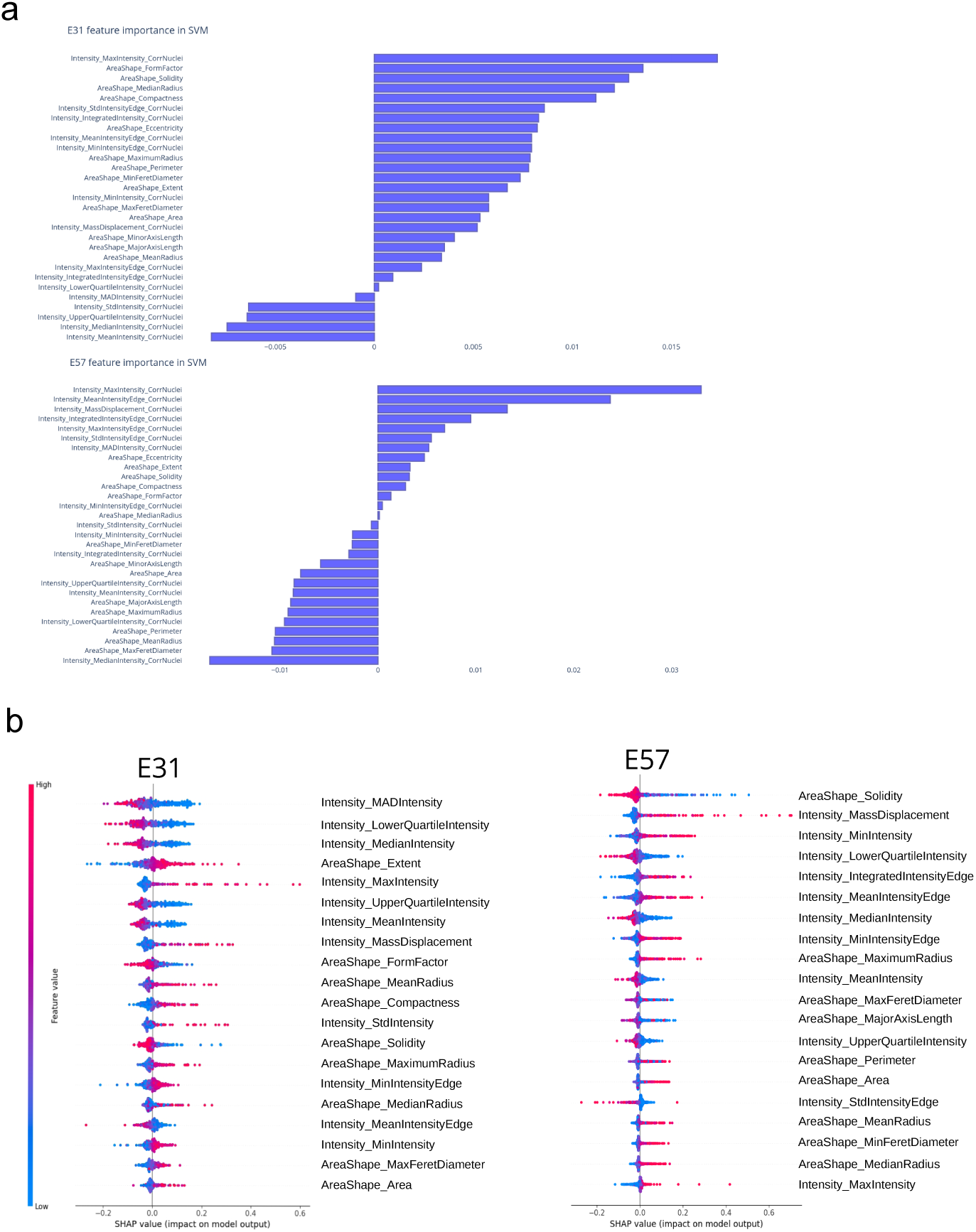
a) Importance of features in the SVM model for cell lines E31 and E57. b) SHAP values for most important model features for cell lines E31 and E57. Features are ordered by importance, with the most important at the top.

**Figure S4:**
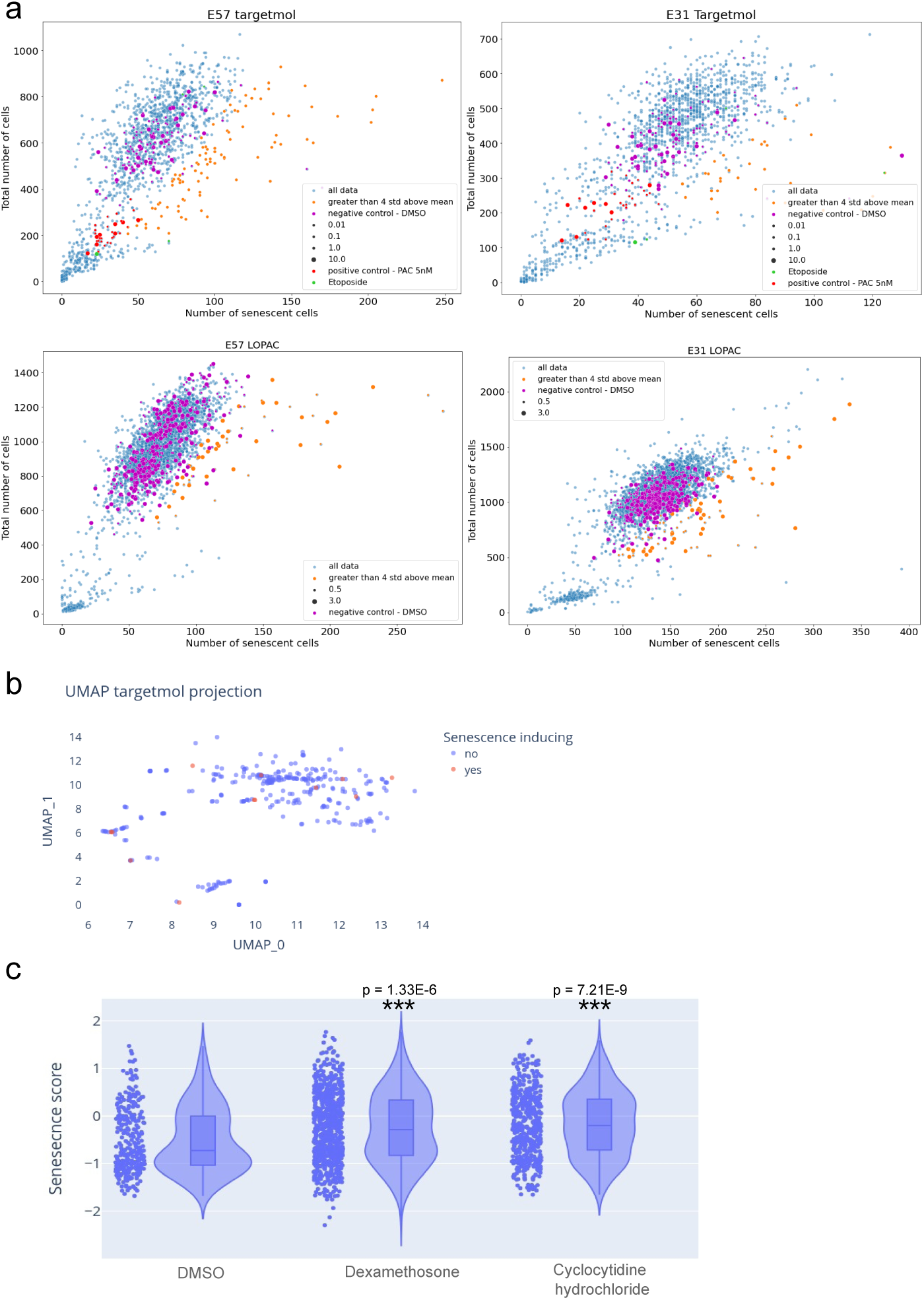
Identification of senescence-inducing compounds. a) The number of cells predicted to be senescent due to the compounds in the Targetmol and LOPAC libraries with DMSO controls, PAC controls, and senescence-inducing compounds highlighted. b) UMAP of the SMILES representation of the compounds in the targetmol library, coloured by senescence-inducing properties. c) Senescence score prediction for 3 compounds from the targetmol library, where analysis started from the raw images of cells treated by the compounds.

**Figure S5:**
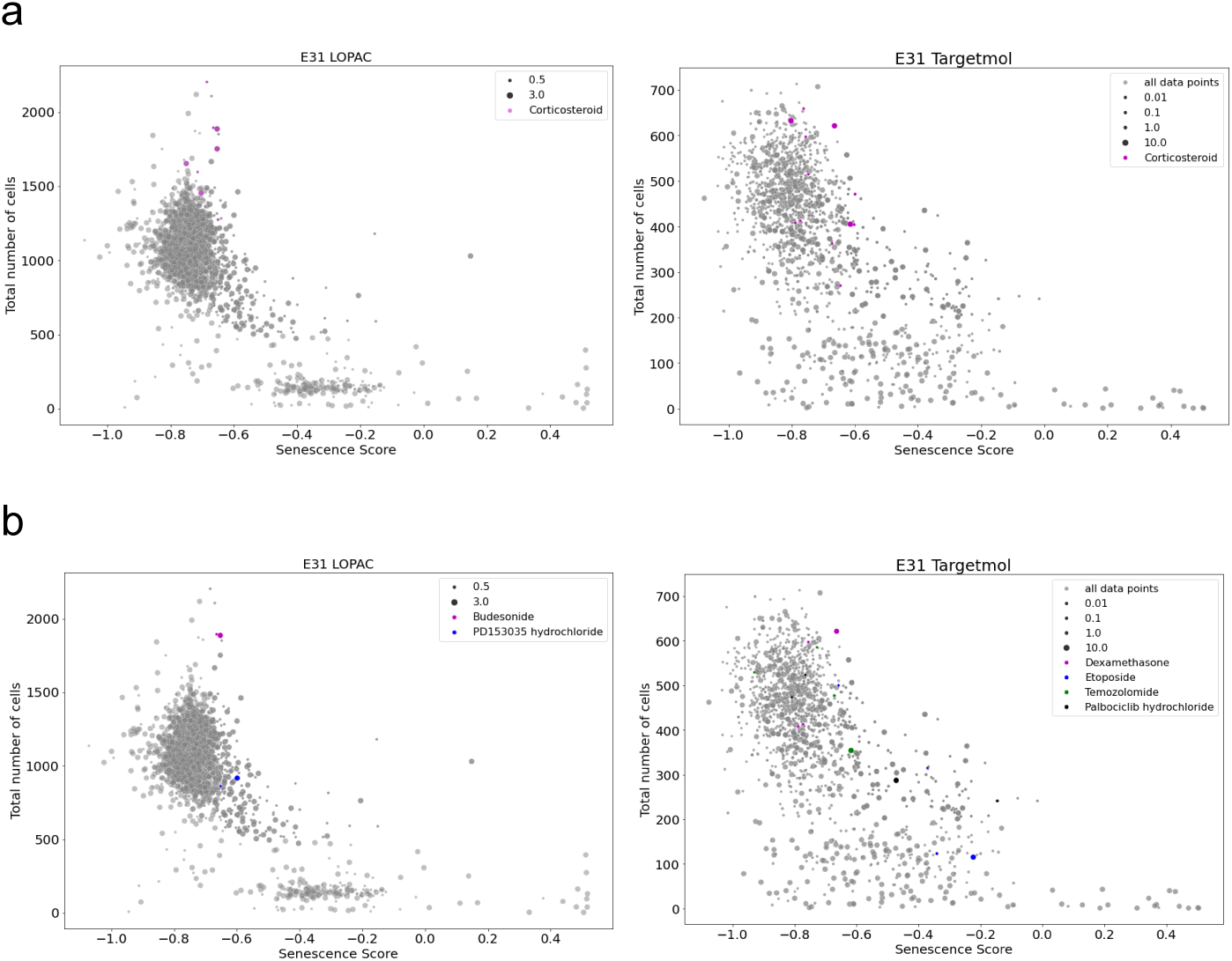
Highlighted compounds of interest in the two high throughput drug screening datasets. a) Glutocorticoids that were found to induce senescence. b) Compounds that we tested in the lab.

**Table S4:** Reduced list of features used in SVM model. Definition taken from [41].

| Feature | Description |
| --- | --- |
| AreaShape_Area | Nuclear area |
| AreaShape_Compactness | The mean squared distance of nuclei's pixels from the centroid divided by the area. A filled circle will have a compactness of 1, with irregular objects or objects with holes having a value greater than 1. |
| AreaShape_Eccentricity | the ratio of the distance between the foci of the ellipse and its major axis length. The value is between 0 and 1. |
| AreaShape_Extent | The area/volume of the object divided by the area/volume of the bounding box. |
| AreaShape_FormFactor | $4*\pi*Area/Perimeter^2$ |
| AreaShape_MajorAxisLength | The length (in pixels) of the major axis of the ellipse. |
| AreaShape_MaxFeretDiameter | The distance between two parallel lines tangent on either side of the object. |
| AreaShape_MaximumRadius | The max distance of any pixel in the object to the closest pixel outside of the object. |
| AreaShape_MeanRadius | The mean distance of any pixel in the object to the closest pixel outside of the object. |
| AreaShape_MedianRadius | The median distance of any pixel in the object to the closest pixel outside of the object. |
| AreaShape_MinFeretDiameter | The distance between two parallel lines tangent on either side of the object. |
| AreaShape_MinorAxisLength | The length (in pixels) of the minor axis of the ellipse. |
| AreaShape_Perimeter | The total number of pixels around the boundary of each region in the image |
| AreaShape_Solidity | The proportion of the pixels in the convex hull that are also in the object, i.e., $ObjectArea/ConvexHullArea$. |
| Intensity_UpperQuartileIntensity | The intensity value of the pixel for which 75% of the pixels in the object have lower values. |
| Intensity_IntegratedIntensityEdge | The sum of the edge pixel intensities of an object. |
| Intensity_IntegratedIntensity | The sum of the pixel intensities within an object. |
| Intensity_LowerQuartileIntensity | The intensity value of the pixel for which 25% of the pixels in the object have lower values. |
| Intensity_MADIntensity | The median absolute deviation (MAD) value of the intensities within the object. |
| Intensity_MassDisplacement | The distance between the centres of gravity in the grey-level representation of the object and the binary representation of the object. |
| Intensity_MaxIntensityEdge | The maximal edge pixel intensity of an object. |
| Intensity_MaxIntensity | The maximal pixel intensity within an object. |
| Intensity_MeanIntensityEdge | The average edge pixel intensity of an object. |
| Intensity_MeanIntensity | The average pixel intensity within an object. |
| Intensity_MedianIntensity | The median intensity value within the object. |
| Intensity_MinIntensityEdge | The minimal edge pixel intensity of an object. |
| Intensity_MinIntensity | The minimal pixel intensity within an object. |
| Intensity_StdIntensityEdge | The standard deviation of the edge pixel intensities of an object. |
| Intensity_StdIntensity | The standard deviation of the pixel intensities within an object. |

